# MPXV Clade IIb virus infection in mice leads to prolonged viral replication, macrophage infiltration, and decreased spermatogenesis in the testes

**DOI:** 10.64898/2025.12.16.694558

**Authors:** Cynthia L. Swan, Gustavo Sganzerla Martinez, Mia Madel Alfajaro, Angela L. Rasmussen, David J. Kelvin, David Evans, Jason Kindrachuk, Ryan S. Noyce, Alyson A. Kelvin

## Abstract

Mpox (formerly monkeypox) is caused by monkeypox virus (MPXV) and has prompted two recent global health emergencies. Clade IIb MPXV, a recently recognized subclade, has been associated with oral and genital lesions and transmission among men who have sex with men (MSM); however, mechanisms of genital pathogenesis and sexual transmission are not understood. We investigated several routes of MPXV Clade IIb virus infection (intranasal, oral, anal, and intraperitoneal) and found high and prolonged viral titres in the testes after IP inoculation still detectable at 21 days. The testes had significant changes to tissue architecture including loss of spermatogenesis, disorganization of spermatozoa, loss of Leydig cells, and breakdown of the seminiferous tubule membranes. Viral antigen positive cells were present in the interstitial spaces between the seminiferous tubules with macrophage infiltration also evident. This work provides insights into potential of sexual transmission for MPXV viruses as well as mechanisms of Mpox disease which may significantly impact long-term fertility or sex organ health of infected males.

**Author Summary:** Mpox (formerly monkeypox) is a painful disease similar to smallpox caused by the monkeypox virus (MPXV) which is an emerging/re-emerging virus. Recently the WHO has declared two global health emergencies due to human-to-human transmission of MPXV. Clade IIb MPXV is a newly recognized subclade that has been associated with genital lesions and sexual transmission/contact; however, how the virus causes disease in sex-organs or how it is passed from person-to-person is not understood. We investigated several routes of MPXV Clade IIb inoculation and found IP inoculation in male mice led to high and prolonged viral titres in testes. Additionally, the testes were significantly affected having loss of sperm and cellular organization. Evidence of inflammatory cell infiltration, specifically, macrophages, are suspected to be the cause of testes specific disease. This work provides insights into potential of sexual transmission for MPXV viruses as well as mechanisms of Mpox disease.

## Introduction

Mpox is a zoonotic disease caused by monkeypox virus (MPXV), an enveloped double-stranded DNA virus with a genome size of ∼197 kb which was first shown to infect humans in 1970 ^1–7^. MPXV (*Orthopoxvirus monkeypox*) is a member of the Orthopoxvirus genus within the family *Poxviridae*. MPXV is divided into two phylogenetically distinct lineages, Clade I (formerly the Congo Basin Clade) ^8^ and Clade II (formerly the West African Clade), which have a 95% nucleotide identity ^1,9,10^. Human infection with MPXV have classically been characterized by a prolonged painful disease often including skin lesions with the disease lasting between 2 and 4 weeks ^11^. Common symptoms of MPXV infection in humans include fever, swollen lymph nodes and a painful rash with lesions that historically were associated with spread across the body in centrifugal distribution ^12^. Historically, Clade I MPXV has been associated with higher case fatality rates compared to infections with Clade II viruses, although the context-dependency of this distinction should be considered ^13^. Previously, human mpox cases have been localized primarily to tropical forested regions of West and Central Africa where the virus is endemic with the majority of infections due to zoonotic contacts with infected animals and limited secondary transmission events ^6,13–15^.

Human mpox outbreaks were historically considered sporadic until the late 2010s, when the number of cases in endemic areas as well as the number of travel related cases began to significantly increase or the identification of cases increased ^16–18^. Recently, two new subclades of MPXVs, Clade Ib and Clade IIb, have been identified which are associated with increased human-to-human transmission and their circulation has prompted two Public Health Emergencies of International Concern (PHEIC) in 2022 and 2024 ^19^. Clade IIb, specifically, was first characterized in July 2022 although evidence suggests it had been circulating in humans for several years prior^20^. Moreover, MPXV Clade IIb appears to be more readily transmissible among humans, which led to local community transmission as well as travel related cases in an unprecedented number of non-endemic countries ^7,21,22^. As of July 2025, more than 150,000 confirmed and probable mpox cases have been reported across 139 countries, with over 410 reported MPXV associated deaths ^23,24^. Moreover, the human-to-human transmission of the 2022 Clade IIb outbreak was linked to close contact, including sexual contact with the majority of reported cases being male and in the demographic of men who have sex with men ^6,25–27^. Although mpox was historically characterized by lesions distributed throughout the body, Clade IIb infections frequently present with lesions localized on the genital and in the oral mucosa ^25,26,28–30^. Studies have demonstrated that viral nucleic acid can be detected in the semen of people who have tested positive for MPXV Clade IIb virus ^31–37^. Although close contact is considered the primary confirmed route for human-to-human transmission, the role of sexual contact or sexual intercourse in transmission has not been thoroughly studied.

Pre-clinical animal models are important tools for studying viral pathogenesis, modes of transmission, mechanism of host immunity, and evaluating the efficacy of vaccines and antiviral therapies. Previous studies have shown that classic small rodent models, such as BALB/c and C57Bl/6 mice, hamsters, and guinea pigs, are not susceptible to MPXV infection ^38,39^. Consequently, several large animal or non-traditional animal models have been used to understand MPXV infection and pathogenesis, including non-human primates, blacktail prairie dogs, squirrels, the African dormouse, the African rat *Mastomys natalensis*, and rabbits ^23,39–47^. However, these models are not easily leveraged for preclinical studies due to ethical considerations, limited animal availability, high costs, and the lack of specifies specific reagents. CAST/EiJ (CAST) mice are a wildtype mouse strain that are susceptible to MPXV infection due to a natural defect in interferon-gamma (IFN-γ) signaling. They have previously been used to study MPXV Clade Ia and IIa viruses through intranasal, intraperitoneal and intradermal routes of inoculation ^44,48–52^. However, studies investigating the pathogenesis, modes of transmission, and exposure routes relevant to sexual contact with Clade IIb viruses in preclinical models are limited.

Here, we investigated four routes of inoculation (intranasal, intraperitoneal, intra-anal, and oral) relevant for the demographic case clusters of the recently described MPXV Clade IIb viruses using the CAST/EiJ mouse model. An extensive analysis of viral tropism and pathogenesis was conducted. For inoculations we used a Canadian MPXV Clade IIb viral isolate. Our results showed systemic MPXV infection with the highest live viral load and viral DNA in the testes associated with macrophage infiltration and decreased spermatogenesis following intraperitoneal inoculation.

## Results

### Intraperitoneal inoculation causes systemic MPXV Clade IIb virus infection with highest levels of virus in the testes

When MPXV Clade IIb viruses were first identified, it was not clear how the virus was efficiently transmitted in humans. To gain insight into potential exposure routes influencing infection susceptibility, we explored inoculation routes using the CAST/EiJ model due to its readily available status as a preclinical model. We selected the intranasal and the intraperitoneal routes based on previous studies demonstrating robust infection of Clade Ia viruses in CAST/EiJ mice ^39,41,50–53^. For our studies we used the MPXV Clade IIb, lineage B.1.4 strain which was isolated from a patient in Canada during the 2022 MPXV outbreak. Only male mice were used to reflect the demographic pattern of Clade IIb infections, which predominantly affect men ^31,34^.

Intranasal inoculation of CAST/EiJ mice at two infectious doses of MPXV Clade IIb virus (1×10^4^ PFU/mouse or 1×10^5^ PFU/mouse) did not lead to weight loss or signs of disease (Supplementary Figure 1B) compared to mock inoculated control mice. Moreover, there was no evidence of robust levels of viral DNA in the tissues (nasal turbinates, lungs, hearts, mediastinal lymph nodes, kidneys, livers, spleens, small intestines, large intestines, and brain) of inoculated animals (Supplementary Figure 1C). Collectively, these results suggested that the intranasal route of inoculation with MPXV Clade IIb virus in CAST/EiJ mice was insufficient to induce a consistent infection and disease.

We next investigated male CAST/EiJ mice intraperitoneally inoculated with 10^4^ or 10^5^ PFU virus per animal. A mock inoculated group was also included as control. Tissues were sampled on Day 6, 12, and 21 (Figure 1A) and clinical signs were monitored daily with animal weights were recorded every other day. Monitoring over the time course indicated that body weight did not decrease and the inoculated animals did not show signs of disease (Figure 1B). However, qPCR analysis of viral DNA in tissues revealed that IP inoculation with both low (10^4^ PFU/mouse) and high (10^5^ PFU/mouse) doses of virus consistently produced high levels of viral DNA in all of the tissues studied except for the brain (Supplementary Figure 2, Figure 1) suggesting systemic virus distribution. Analysis of tissues from animals inoculated by IP injection with 10^5^ PFU indicated that viral DNA levels peaked at Day 6 for all tissues analyzed (Figure 1 C-F). Viral DNA levels are graphed according to anatomical regions (Head and Upper Thorax; Thorax; Gut; and Urogenital regions) of the tissue analyzed. The lung, heart and MLN (Upper Thorax) (Figure 1C), kidney, liver and spleen (Lower Thorax) (Figure 1D), small and large intestine (Gut) (Figure 1E) and testes, penis, and bladder (Urogenital) (Figure 1F) all had positive C_T_ values (those above the LL (Lower Limit) of detection) starting on Day 2 post inoculation. By Day 12 post inoculation, the heart, brain, MLN, kidney, liver, spleen and penis viral DNA levels dropped below or equal to the LL. Tissues with C_T_ values that persisted to Day 21 included the lungs, small intestine, large intestine, bladder, and testes. The highest levels of viral DNA were detected in the bladder and testes over the entire time course. At peak infection on Day 6, the average C_T_ value for the testes, bladder and large intestine were 15.0, 18.4 and 19.4, respectively, while C_T_ values for all other tissues ranged from 34 to 20. Trends of C_T_ values per tissue over time for mice inoculated with 10^5^ PFU are shown in Supplementary Figure 2. Similar trends were observed in viral DNA levels in the 10^4^ PFU inoculated animals (Supplementary Figure 3) again with prolonged high levels of viral DNA being detected on Day 21 post inoculation in the testes and bladder.

**Figure 1.**
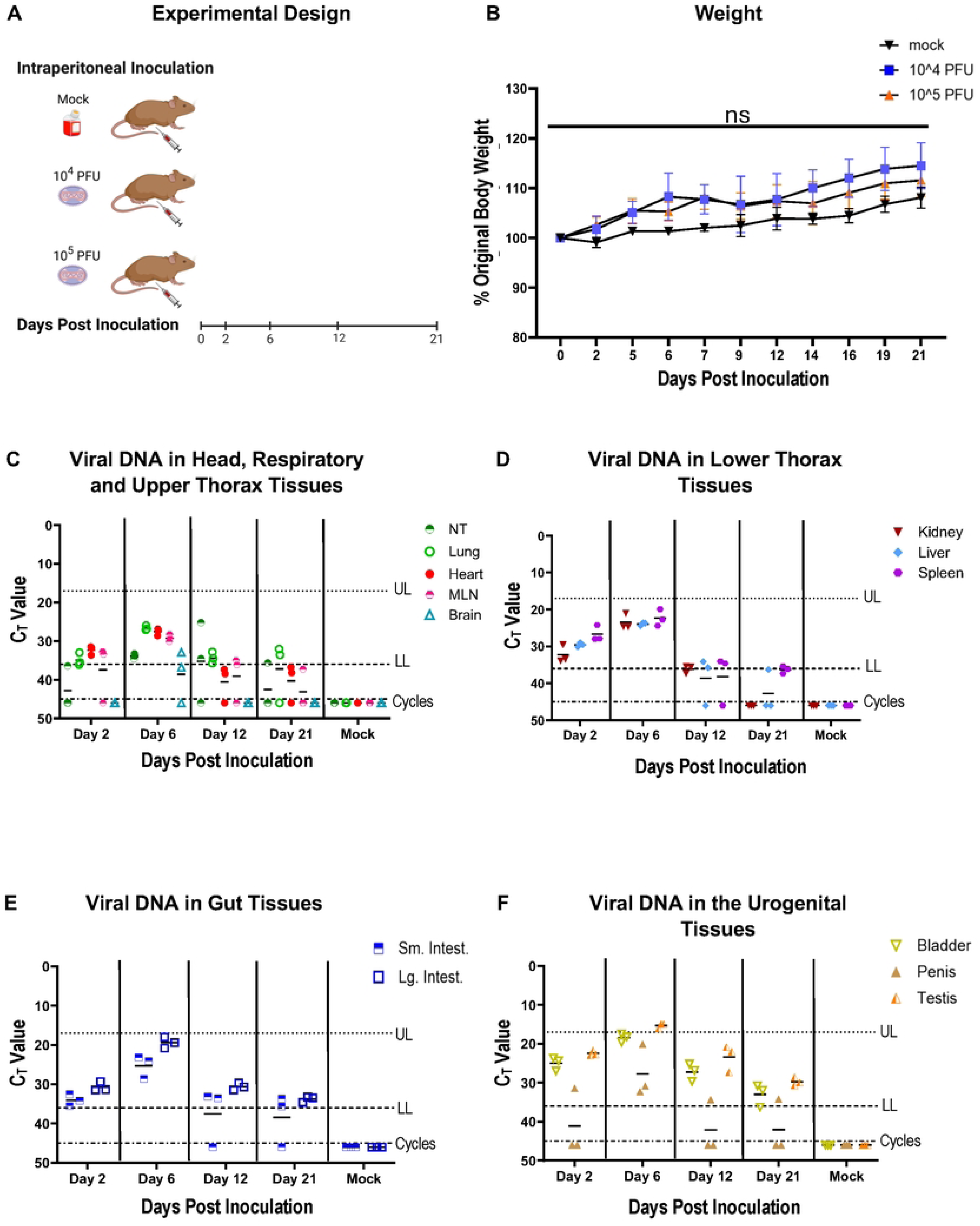
Intraperitoneal inoculation with MPXV Clade IIb results in systemic tissue infection, highest viral DNA in the testes and no weight loss in CAST/EiJ mice. Male mice were inoculated with either 1×10^4^ PFU or 1×10^5^ PFU MPXV Clade IIb per animal or media (mock) through the intraperitoneal route of infection. A group inoculated with media through the intraperitoneal route was used as control. Tissues samples were collected for analysis at 2-, 6-, 12-, and 21-days post inoculation (n=3 per group per time point). The experimental schematic is shown (A). Percent original body weight per group over the 21-day time course did not indicate weight loss (n≥8 per group) (B). Quantitative PCR was used to determine MPXV Clade IIb DNA levels in each tissue. The resulting critical threshold (C_T_) values from mice infected with 1×10^5^ PFU MPXV Clade IIb are shown in graphs (C) to (F). Data was grouped into body regions: head, respiratory and upper thorax tissues (C), the lower thorax (D), the gut tissues (E) and the urogenital tract are shown (F). The lower limit (LL) and upper limit (UP) of the assay range and total cycles used (45) are denoted by dashed lines determined as previously described. All tissues with an undetermined C_T_ value were assigned a value of 46 for graphing purposes. Significant differences in weight loss were determined by Tukeys pairwise comparison test. ns indicates no significant difference was found between the three experimental groups for percent change of body weight. The experimental design schematic was created in https://BioRender.com.

As these analyses suggested that MPXV can replicate in different tissues, plaque assays were used as a confirmation of viral replication. We focused on tissues collected on Day 6 following 10^5^ PFU IP inoculations for quantifying viral titer in tissue by plaque assay (Figure 2). Plaques were identified from most tissues analyzed demonstrating the presence of infectious virus (Figure 2A, right panel). Similar trends were observed between the PFU/mg for each tissue with the viral DNA quantification for that tissue. The highest levels of infectious virus were detected in the testes and bladder, with titres between 10^3.5^ and 10^4^ PFU/mg tissue supporting the viral DNA results. One of the three penile tissues also had similar viral titres as the testes and bladder. The intestines, kidney, spleen, and MLN also had robust viral titres ∼ 10^2^ PFU/mg. Of note, we were not able to accurately quantify virus in liver samples possibly due to toxicity of the tissue on cells in culture during the plaque assay; however, at lower dilutions of the tissue homogenate, plaques were observed (potentially suggesting the liver is also a site for virus replication).

**Figure 2.**
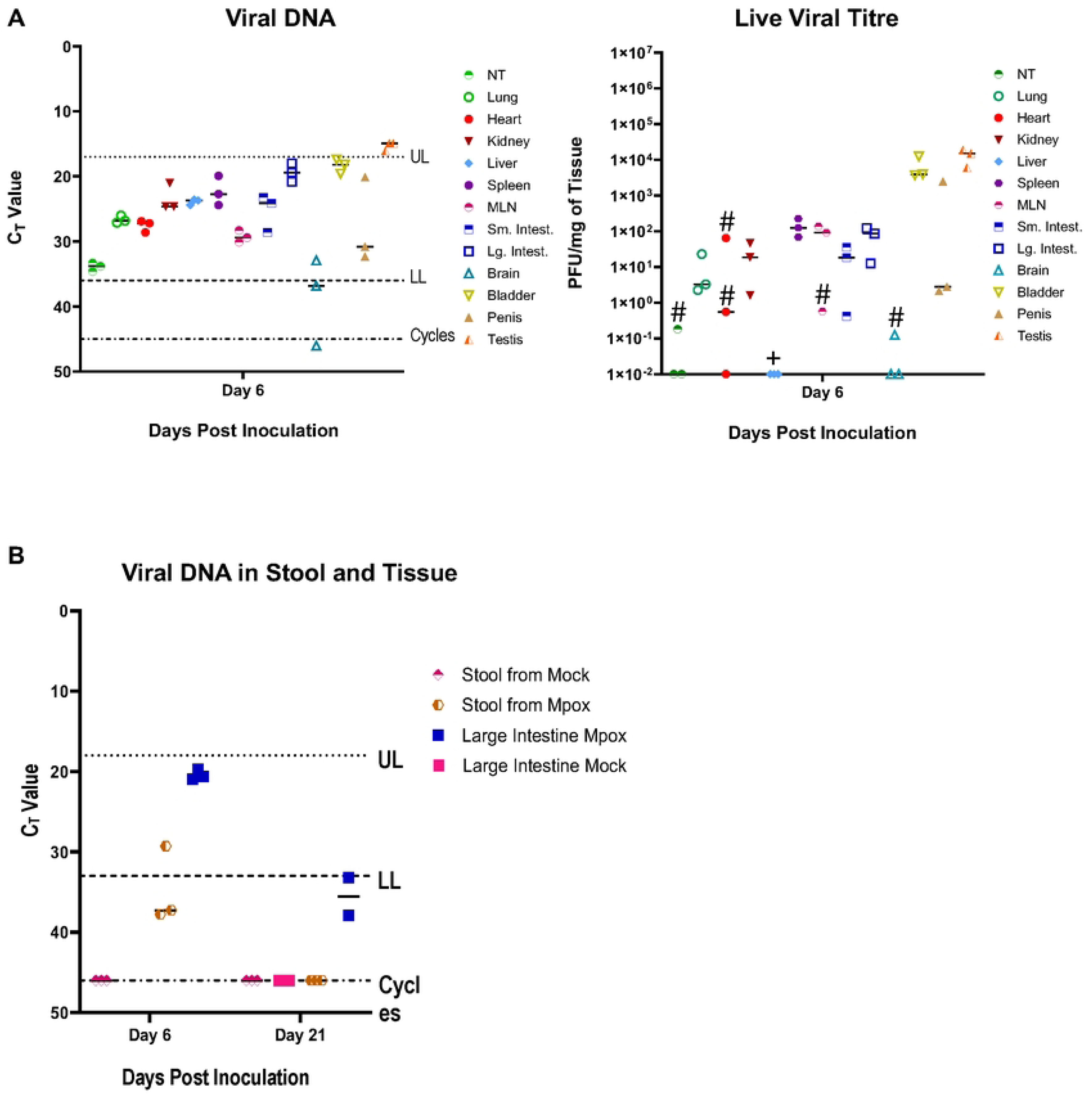
Highest viral titres in testes with high titers in the large intestines and bladder of CAST/EiJ mice inoculated intraperitoneally with 1×10^5^ PFU MPXV Clade IIb. Tissues were collected from male mice (n=3 per group per time point) 6 days post inoculation with 1×10^5^ PFU MPXV Clade IIb per animal. Tissue homogenates were analyzed by Taqman qPCR for MPXV DNA (A, left panel) and by plaque assay to determine infectious virus titres (A, right panel). The graphs are placed side-by-side for comparison. (B) qPCR was used to detect MPXV Clade IIb viral DNA in the stool of mice with high C_T_ (critical threshold) levels of viral DNA in corresponding large intestines after intraperitoneal inoculation with 1×10^5^ PFU MPXV Clade IIb. Samples were collected from 3 mice per group per timepoint. DNA content of tissues is shown as raw C_T_ values from the PCR reactions with the LL and UP as described previously. All tissues with an undetermined C_T_ value were assigned a value of 46 on the graph. Infectious viral titres in tissues were calculated as PFU/mL of tissue homogenate by plaque assay. # = a single plaque was observed out of all replicates of the four dilutions plated. A single plaque was observed for two biological replicates of heart tissues and one replicate of NT, MLN, and brain tissue. The + symbol indicates that, while some plaques were observed in the liver tissues, the amount of cell death due to tissue toxicity of the cells made the calculation of total plaques unreliable and therefore all values were set to 0.

Since high levels of infectious virus and viral DNA were present in the large intestines, we also used qPCR to quantify the viral DNA in stool collected from mock and 10^5^ PFU IP inoculated mice at Day 6 and 21 post inoculation. This analysis showed a low level of viral DNA was present in stool on Day 6 which is shown in comparison to the levels of the large intestine of the corresponding animal (Figure 2B). By Day 21, viral DNA was still detected at low levels in the large intestine, while it was undetectable in the stool.

### Marked tissue disorganization, viral antigen presence, and macrophage infiltration in the testes of mice infected with MPXV Clade IIb virus by intraperitoneal inoculation

Tissues collected from mice inoculated at 10^5^ PFU IP were investigated by histopathology and evaluated by a board-certified pathologist (Figure 3; Supplementary Figure 4 and 5). No major differences in architecture were observed between infected and non-infected mice for most of the tissues observed (heart, lung, kidney, liver, spleen, small intestine, large intestine, brain) (Supplementary Figure 4). Small inflammatory lesions (score of 1 or 2) were noted on the outer serosa of the large intestine tissues from 3 out of 6 infected animals which were attributed to a viral infection. However, the tissue of the urogenital tract including testes, bladder, and penis, from Day 6 and Day 21 had moderate (score 2 to 3) to severe (score 3 to 4+) scores (Figure 3 and Supplementary Figure 5). Tissues from the urogenital tract of mock inoculated animals had no lesions or hallmarks of infection and inflammation present (Figure 3 A, B, C, and D (Testes); Supplementary Figure 5 G, H, I (Bladder) and P, Q, R (Penis)). In the testes at Day 6 post inoculation, H&E analysis revealed a prominent loss of spermatogenic cells with concomitant absence of spermatozoa and disorganization of spermatogenic cells (Figure 3; loss of spermatozoa is highlighted by red plus sign). Moreover, on Day 6 (Figure 3 E, F, G, and H) and Day 21 Figure 3 I, J, K, and L) post MPXV Clade IIb inoculation, there was loss of Leydig cell proliferation (black asterisks) and increased interstitial space between the seminiferous tubules where green arrows mark interstitial space between in the seminiferous tubules of control and inoculated mice. The lumen of seminiferous tubules (ST) became vacuous on Day 6 post inoculation compared to the mock (the lumen of inoculated and control seminiferous tubules are denoted by black arrows in Figure 3). In the infected animals, there was increased infiltrating leukocytes in the interstitial space and the surrounding tissues of the testes in inoculated animals (Figure 3; blue stars indicate tissue surrounding the testes). Cell infiltration was noted in the bladder and penis of infected animals (Supplementary Figure 5A-F and J-O). Immunohistochemical analysis for viral protein antigen revealed significant staining in the testes and bladder on Day 6 (Figure 4 E, F, G and H; Supplementary Figure 6 A, B, and C). In the testes and bladder on Day 6 post inoculation, there was infiltration of viral antigen positive inflammatory cells suspected to be macrophages (Figure 4 denoted by yellow outlined black arrows). Viral antigen positive cells decreased in the testes by Day 21 (Figure 4 I, J, K, and L). To investigate if the infiltrating cells in the testes and bladder were macrophages, we stained these tissues with the macrophage antibody IBA1 (Figure 5 and Supplementary Figure 7). Results indicated that macrophages (Figure 5, black arrows) were similarly distributed in the testes on Day 6 as seen with the antigen positive cells in the same tissue. IBA1 positive cells were found in the interstitial space between seminiferous tubules and the tissue surrounding the testes (Figure 5 G and K, black arrows). In the bladder (Supplementary Figure 7), presumptive macrophages were found in the serosa, surrounding smooth muscle cells, and the Lamina propria.

**Figure 3.**
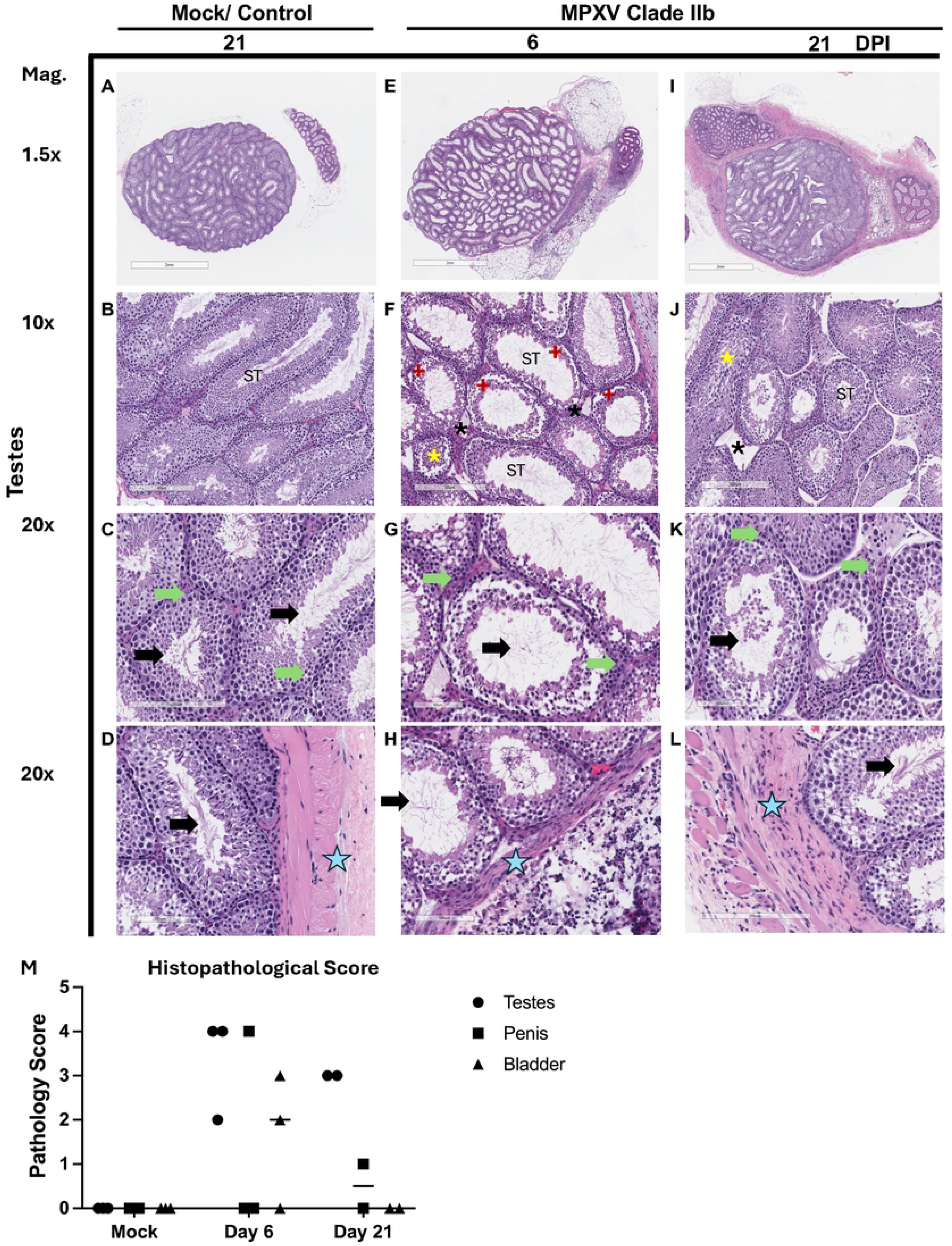
Decreased spermatogenesis, increased leukocyte infiltration, and breakdown of the membranes of the seminiferous tubules in testes following MPXV Clade IIb infection. Male mice were inoculated interperitoneally with 1×10^5^ PFU MPXV Clade IIb or media (mock) per animal as described previously. Tissues were collected on Day 6 (E, F, G, and H; n=3) and 21 (I, J, K, and L; n=2) post inoculation with MPXV Clade IIB or after mock inoculation (Day 21 post mock inoculation) (A, B, C, and D). Tissue samples were fixed in 10% formalin, paraffin embedded, mounted, and stained with a Hematoxylin and Eosin stain. Tissue sections from testes are shown at 1.5x, 10x and 20x magnification. Specific areas of the testes and pathological features are denoted: ST = Seminiferous Tubules; * = loss of Leydig cell proliferation; red star = degeneration of spermatogenic cells with absence of spermatozoa; yellow asterisk = disarrangement of spermatogenic cells were observed; green arrows = interstitial space between seminiferous tubules; black arrows = lumen of seminiferous tubules; and blue stars = tissue surrounding the testes. Scale bars, 2 mm for figures at 1.5x magnification, 300 µm for 10x magnification and 100 µm or 200 µm for 20 x magnification.

**Figure 4.**
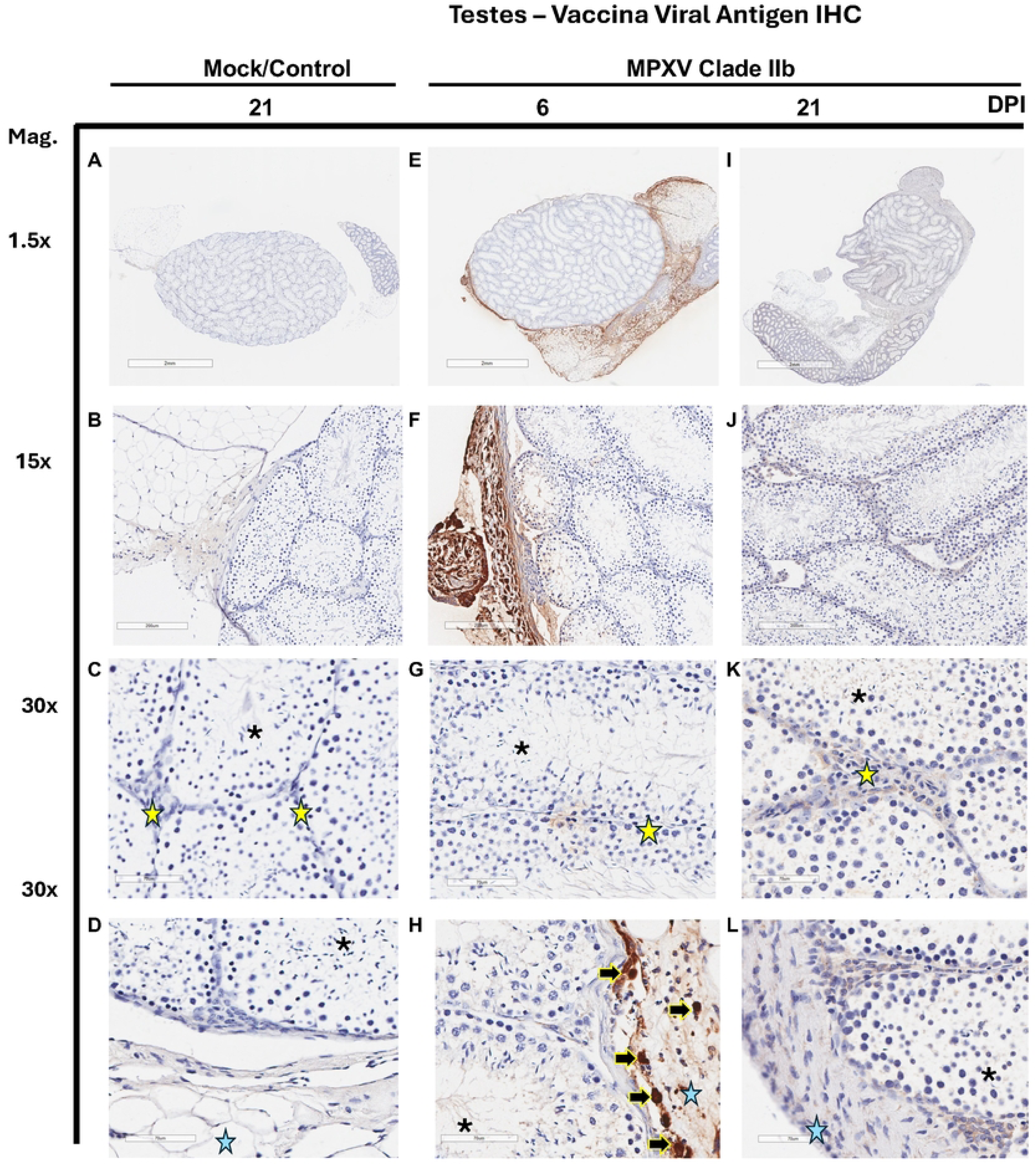
Poxvirus antigen detected in testes of CAST/EiJ mice 6 and 21 days after IP inoculation with MPXV Clade IIb. Testes tissue samples collected on Day 6 and 21 post virus inoculation or on Day 21 for the Mock inoculated animals. Tissues were fixed in 10% formalin and paraffin embedded. Viral antigen (brown) was detected in testes by IHC using an anti-Vaccinia virus antibody (abcam 35219). Representative tissue sections of testes from mice Mock inoculated control (Day 21) (A, B, C, and D; n=3), 6 days post inoculation (E, F, G, and H; n=3) and 21 days post inoculation (I, J, K, and L; n=2) are shown. Specific areas of the testes and pathological features are denoted on the 30x panels: yellow outlined black arrows = viral antigen positive cells; yellow stars = interstitial space between seminiferous tubules; black asterisks = lumen of seminiferous tubules; and blue stars = tissue surrounding the testes. Scale bars at 2 mm for images at 1.5x magnification, 200 µm for 15x magnification and 70 µm for 30x magnification.

**Figure 5.**
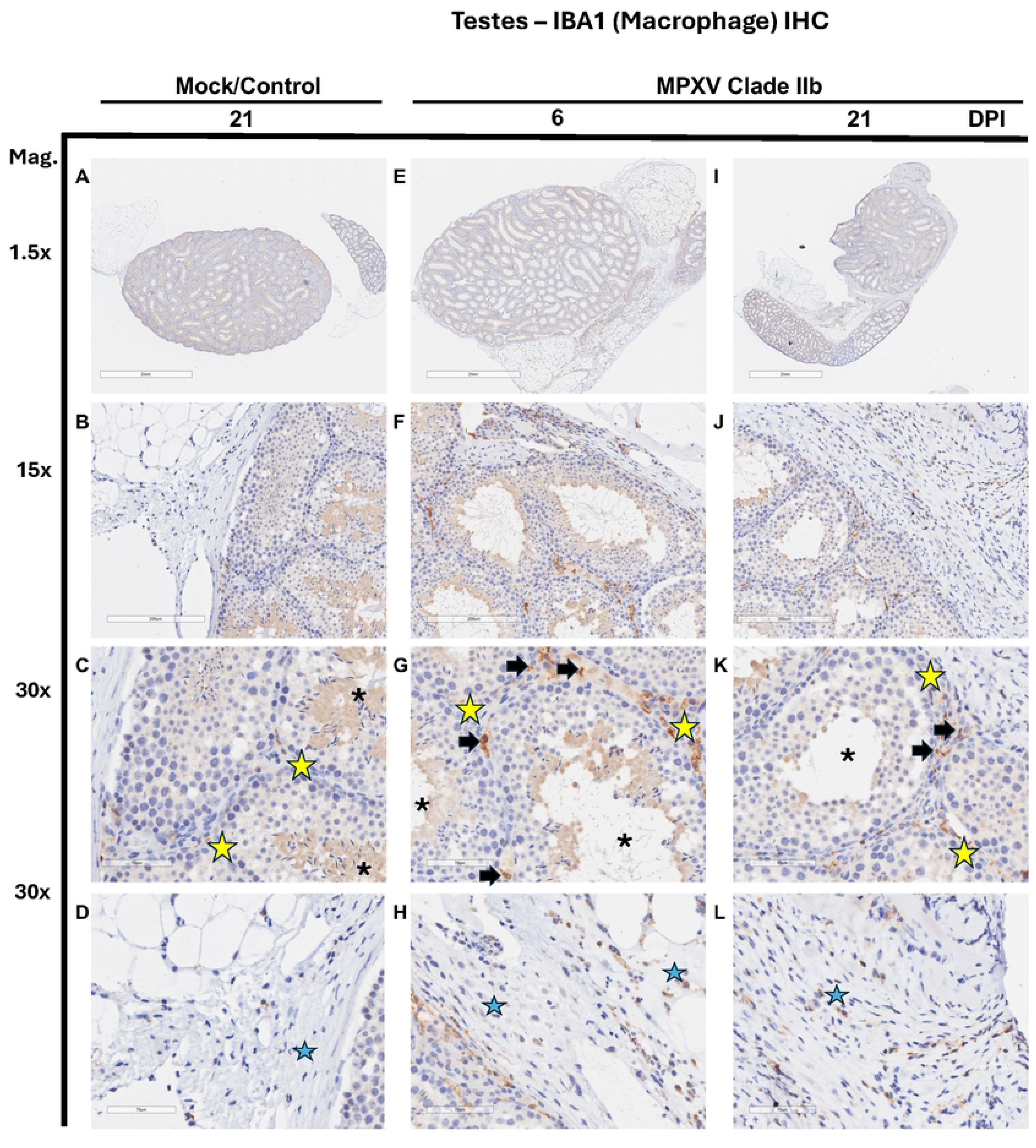
An increase in macrophage IBA1 antigen positive cells in testes of CAST/EiJ mice 6 and 21 days after IP inoculation with MPXV Clade IIb. Tissues from Mock IP inoculated and 1×10^5^ PFU MPXV Clade IIb inoculated mice were fixed in 10% formalin and paraffin embedded. IBA1 antigen (brown) was detected in testes by IHC using rabbit anti-IBA1 antibody (ThermoFisher PA5-27436). Representative tissue sections of testes from mice Mock inoculated controls (Day 21) (A, B, C, and D; n=3), Day 6 post inoculation with MPXV Clade IIb (E, F, G, and H; n=3) and Day 21 post inoculation with MPXV Clade IIb (I, J, K, and L; n=2) are shown. Specific areas of the testes and pathological features are denoted on the 30x panels: black arrows = IBA1 positive cells (presumptive macrophages); yellow stars = the interstitial space between seminiferous tubules; black asterisks in the lumen of seminiferous tubules; and blue stars indicating the tissue surrounding the testes. Scale bars at 2 mm for images at 1.5x magnification, 200 µm for 15x magnification and 70 µm for 30x magnification.

### Anal and oral routes of inoculation do not result in infection

Above, we showed high titres of live MPXV Clade IIb virus in the large intestine and in the urogenital tract following IP inoculation. Several human studies have suggested that Clade IIb virus can be transmitted by sexual contact, with a high proportion of global infections among males, and predominantly those that identify as GBMSM ^25,26^. Therefore, we assessed infection susceptibility following MPXV Clade IIb exposure in the anal and oral cavities as a proxy for investigating sexual contact-mediated virus transmission. Here, we inoculated male CAST/EiJ mice with 10^5^ PFU of MPXV Clade IIb virus through the anal or oral route. Mock anal inoculation with media was used as a study control. We observed clinical signs and collected tissues through a 12-day study period with tissues collected for viral load pathology assessment on Day 2, 6, and 12 post inoculation (Figure 6A). We chose a shorter study period since the robust infections of IP inoculations peaked by Day 12 post inoculation in our previous study. Following anal and oral inoculations, minimal weight change was observed in mice over the 12-day observation period (Figure 6B). Evaluation of viral DNA in tissues revealed inconsistent low levels of virus across various tissue types and time point replicates with only one of the three replicate animals being positive for viral DNA per tissue per group. As well as these levels being inconsistent among replicates, the viral DNA levels were also below the limit of detection (Figure 6C). No tissues were positive for viral DNA C_T_ values above or below the limit of detection from any mouse sampled in the mock control group.

**Figure 6.**
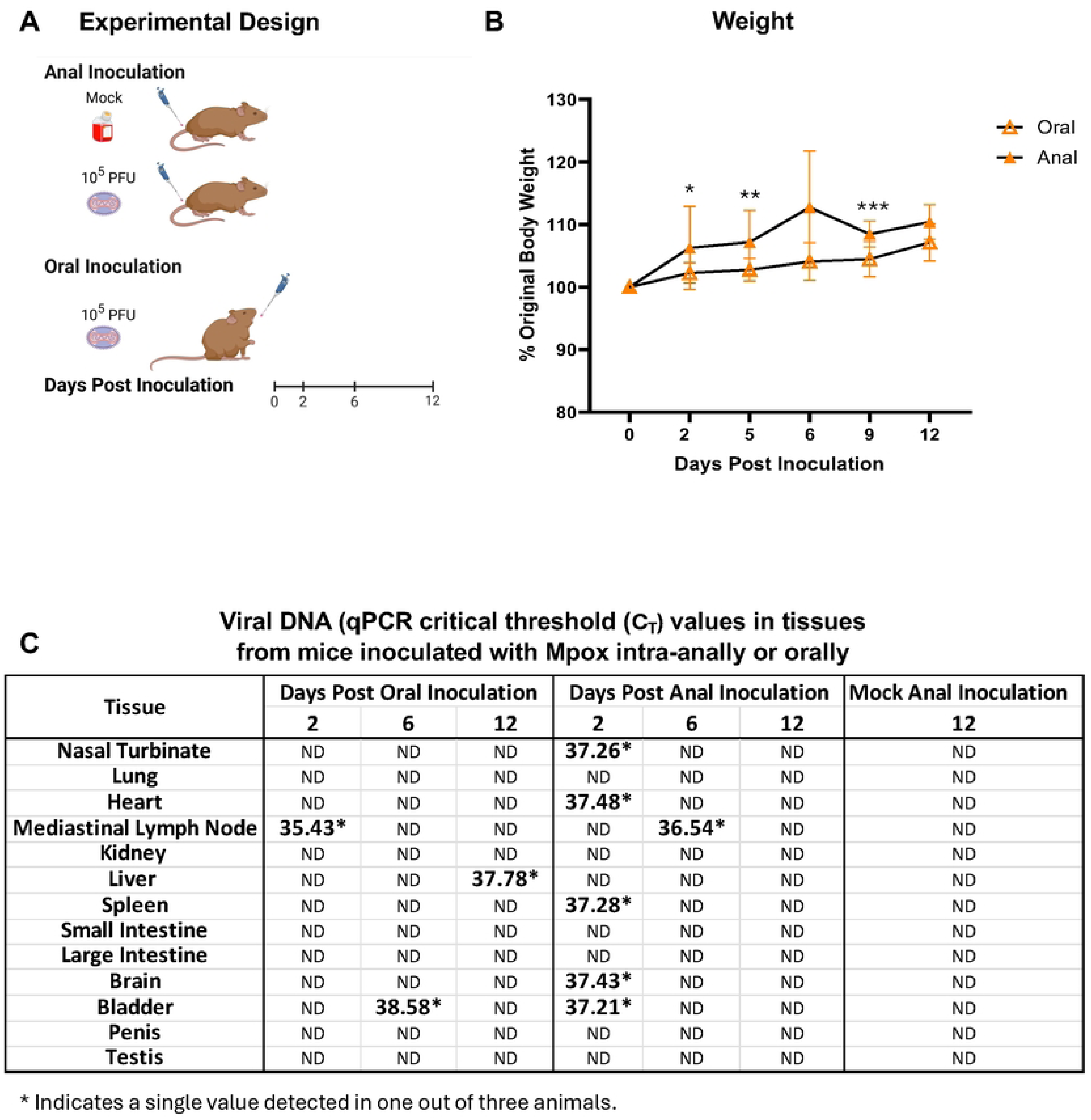
Anal and oral inoculation with MPXV Clade IIb does not result in infection or weight loss in CAST/EiJ mice. Male mice were inoculated with 1×10^5^ PFU MPXV Clade IIb per animal through either an intra-anal or oral route or media inoculated by intra-anal inoculation as a mock control. Tissues samples were collected for analysis at Day 2, 6, and 12 post inoculation (n=3 per group per time point). The experimental schematic is shown (A). Percent original body weight is shown over the time course per group (n≥9 per group) (B). Taqman quantitative PCR was used to determine MPXV DNA levels in tissues (C). DNA was undetectable (ND) or had a very high C_T_ in all tissues (C). * denotes significant differences between anal inoculated and oral inoculated animals where * p ≤ 0.05; ** indicates p ≤ 0.01, and *** indicates p ≤ 0.001. The * symbol in (C) denotes that only one of three biological replicate tissue samples had a detectable C_T_ value while the remaining two samples had undetectable levels of DNA. The experimental design diagram was created in https://BioRender.com.

## Discussion

MPXV are a significant health threat to humans with two public health emergencies of international concern declared by the World Health Organization in 2022 and 2024 ^54,55^. However, the clinical pathogenesis, modes of transmission, evolution, and primary reservoirs of the MPXV viruses including the nuances of clades and subclades are poorly defined. Here, we investigated the susceptibility, viral tropism, pathogenesis, and exposure routes of Clade IIb using the CAST/EiJ model. We found enhanced and prolonged MPXV Clade IIb viral infection in the male urogenital tract as well as high virus replication in the large intestine following IP inoculation. Importantly, marked pathological features were identified in the testes of infected mice which included increased size of the lumen, disorganized alignment of Sertoli cells and reduced sperm in the seminiferous tubule as well as breakdown of the seminiferous tubule membrane itself. Additionally, the presence of viral antigen-positive cells within the testes, along with macrophage infiltration into the same area, suggests that macrophages may contribute to testicular pathogenesis during MPXV Clade IIb virus infection. The development of a robust animal model is critical to facilitate studies of the pathogenesis and infection characteristics of emerging MPXVs as well as to test new mpox vaccines and antiviral drugs. Our work supports evidence from human clinical studies that MPXV Clade IIb viruses may transmit through sexual contact and/or intercourse and provides a role specifically for the testes in this transmission.

Historically, MPXV human-to-human transmission was thought to be infrequent with most infections related to zoonotic contacts and limited secondary transmissions ^6,14,56^. In 2022, MPXV Clade IIb underwent rapid global expansion among non-endemic regions ^16^, where initial clinical reports from mpox patients suggested that infections may be due to human-to-human transmission. Moreover, of the first cases related to the 2022 global outbreak, the majority were males in the GBMSM community and clinical presentation included atypical characteristics such as asynchronous lesion development on the genitals, anus, and oral region ^12,16,25,57^. Sexual contact and/or sexual intercourse was a suspected driver of transmission ^30,58–60^ and further supported by MPXV Clade IIb isolation from human semen ^30,31,61–63^. In our study we investigated inoculation routes that may be associated with sexual contact-related transmission to give insight into these clinical data. Interestingly, our data showed high and prolonged Clade IIb viral loads in the testes, penis, bladder, and large intestine following IP inoculation but not following intranasal, oral, or anal inoculations. Although we found high virus levels in the testes and urogenital tract following IP inoculation, we did not observe the development of skin lesions in these anatomical regions or elsewhere on the infected mice despite internal pathological changes in the testes. Lack of lesion development could be due to the study being performed in the mouse model; as previous work has suggested that CAST/EiJ mice are unable to develop skin lesions ^41,52^. In addition to lack of skin lesions, we also found no apparent clinical signs such as weight loss in the mice infected by IP which was similar to findings in other studies with Clade IIb ^50^. In summary, despite the lack of lesions and weight loss, the data we have put forth on the prolonged viral load and pathogenesis in the testes following Clade IIb IP inoculation brings potential insight into the modes of transmission associated with MPXV Clade IIb in humans while representing a preclinical model for future studies of male specific pathogenesis.

As noted above, we did not find evidence of productive infection follow intranasal inoculation although some previous studies with CAST/EiJ mice were successful with Clade IIb intranasal inoculation ^50,51^. Variable outcomes in infection potential may be due to inoculation volumes where larger volumes were associated with productive intranasal infections. Moreover, the commonly used mouse wildtype strains C57Bl/6 and BALB/c mice have recently been investigated for susceptibility following MPXV Clade IIb intranasal inoculation using larger inoculums, despite these strains being originally thought not to be susceptible to MPXV infection ^64,65^. In these recent studies, both mouse strains were infected after intranasal inoculation; however, only the BALB/c strain developed significant weight loss and clinical disease suggesting a model with a clinical readout useful for vaccine and treatment studies. However, neither study examined viral load in the male urogenital tracts following intranasal infection leaving it unknown if an intranasal infection can lead to productive infection within the testes.

The major finding in our study was that testes have high viral loads associated with increased macrophages, decreased spermatogenesis, and disorganization of the seminiferous tubules within infected testicular tissue. Previous studies have included pathogenic analysis in the male urogenital tract as secondary endpoints following MPXV infection, however, using other animal models. Liu and colleagues reported in a retrospective study that MPXV Clade I and IIa virus is present and may persist in the testes of crab-eating macaques identified through histopathological analysis ^66^. In this previous study, viral antigen was identified in the epididymus, prostate gland, interstitial tissue, spermatozoa, inflammatory cells of the seminiferous tubules, and in the epididymal ducts and epithelium. In a pathogenesis study in adult rabbits, high infectious doses of Clade IIb inoculated in the ear led to disorganization of the seminiferous tubules and germ cell alignment as well as reduction of the size of the tubule lumen size and evidence of cellular necrosis suggesting subclinical disease preferentially affecting the male reproductive tissues ^23^. These findings and others in rabbits supported our results that viral DNA and live virus in the testes can be accompanied by tissue pathology ^67^. Interestingly, we also found an increase in viral antigen positive cells as well as infiltration of macrophages in the testes of inoculated mice who also had a marked decrease in sperm suggesting a decline in spermatogenesis. Studies investigating the relationship between testicular macrophages and sperm development have demonstrated that testicular macrophages promote steroidogenesis through regulating testosterone production in Leydig cells, a process that is crucial for promoting sperm development. ^68–70^. Considering the known role of macrophages in testes health, more work is required to delineate potential mechanisms of MPXV pathogenesis in the testes and how infiltrating macrophage imbalances could impact Leydig cell testosterone production and downstream spermatogenesis following infection.

MPXV is a significant public health threat and ongoing disease burden is a concern in the coming years given the identification of ongoing transmission in both endemic and non-endemic settings^71^. In recent years, there has been increasing appreciation for expanded clinical and transmission characteristics for MPXV ^16^. In particular, sexual contact and/or intercourse-mediated transmission and the role that the male reproductive and urogenital tracts play in transmission and disease progression are of focus. We recognize that understanding the mechanisms of MPXV pathogenesis in the testes and the role this may have in viral transmission are significant public health priorities. Moreover, developing therapeutics for viral clearance or resolving pathogenesis may be a key goal for patient support. Here we are the first to show that IP inoculation of CAST/EiJ mice leads to productive systemic infection in all tissues except the brain with the highest viral loads observed in the testes which remained positive for viral antigen until at least Day 21 post inoculation. Pathogenesis in the testes including decreased sperm number was associated with macrophage infiltration. This data is important for understanding transmission and pathogenesis of the newly identified MPXV Clade IIb viruses as well as developing and testing medical countermeasures that can be used during health emergencies and to prevent outbreaks.

## Acknowledgements

The authors would like to thank Dr. Christina Hutson her guidance, expertise, and support in designing these studies. Additionally, the authors thank Drs. Andrew Van Kessel, Magen Francis, Antonio Facciuolo, and Carla Norleen for support of the studies.

Alyson A. Kelvin is supported by grants from the Canadian Institutes of Health Research (CIHR) including the CIHR Team Grant: Building capacity in interdisciplinary research on mpox (monkeypox) and other (re)emerging zoonotic threats to health (202302MZ1-499095-MZT-CCAA-242177) and a CIHR Project Grant: Pandemic Preparedness and Health Emergencies Research (PPE-185810).

## Author Contributions

AAK conceived and designed the experiments. AAK supervised the study. CLS and AAK conducted the experiments. AAK and CLS wrote the paper. AAK, CLS, MMA, GSM, RN, DJK, and ALR analyzed data. DE and JK provided constructive suggestions for the work. AAK, RSN, and DE secured funding or provided materials for the study.

## Materials and Methods

### Ethics statement

Animal work was performed in accordance with the Canadian Council of Animal Care (CCAC) guidelines, AUP number 20220115 by the University Animal Care Committee (UACC) Animal Research Ethics Board at the University of Saskatchewan. All mouse procedures were performed under 5% isoflurane anesthesia.

### Viruses

The MPXV Clade IIb (lineage B.1.4) virus was isolated from a patient in Alberta, Canada during the 2022 MPXV outbreak. This sequence has been deposited in Genbank (accession # OP586622).

The isolate was cultured by Drs. David Evans and Ryan Noyce in the Containment Level 3 (CL3) facility at the University of Alberta (Edmonton, Alberta).

### Cell lines

BSC-40 cells (ATCC CRL-2761, African green monkey kidney epithelial cells) were maintained in Minimum Essential Medium (Gibco cat# 11095-080) containing 5% fetal bovine serum (Wisent Bioproducts (Cat # 090–150), 5 mL of 100x penicillin (10,000 U/mL)/streptomycin (10,000 μg/mL) (Gibco Cat #15140-122), 5 mL of L-Glutamine (100x) (Gibco Cat# 25030-081), 5 mL of MEM non-essential amino acids (100x) (Gibco Cat# 11140-050), and 5 mL of sodium pyruvate (100x) (Gibco Cat# 11360-070). Cells were grown at 37°C in 5% CO_2_.

### Mice

Male CAST/EiJ mice, 8-weeks of age, were purchased from Jackson Laboratories (Bar Harbor, ME, USA (Cat # 000928).

### Virus growth and titration

MPXV Clade IIb virus stocks were grown on BSC-40 cells cultured in viral MEM (vMEM) (MEM supplemented with 2% FBS, 1000 U/mL penicillin, 1000 ug/mL streptomycin, 2 mM L-Glutamine, 1x non-essential amino acids, and 1 mM sodium pyruvate). Inoculated cells were maintained at 37°C, 5% CO_2_ for 72 hours until 80-90% CPE was observed. Cells were collected by scraping, pelleting and resuspending in phosphate buffered saline (PBS). Virus was collected in the supernatant after a series of sonication and freeze thaw cycles. MPXV was purified through a 36% sucrose cushion prior to use in animal trials. Titre of the MPXV stock was determined by plaque forming assays. All work with infectious MPXV for the study was performed in the Vaccine and Infectious Disease Organization’s (VIDO) CL3 facility (InterVac) in Saskatoon, Saskatchewan, Canada.

### Animal infections and tissue collection

Male CAST/EiJ mice were acclimated in cages at VIDO after arrival for one week prior to viral inoculation. Mice were anesthetized with 5% isoflurane prior to any procedures. Intranasal inoculations were performed by introducing 10µL of MPXV at a concentration of either 10^4^ or 10^5^ PFU per animal into a single nare. Intraperitoneal inoculations were performed by injecting 200 µL of MPXV at a concentration of either 10^4^ or 10^5^ PFU per animal. Oral inoculation was performed by pipetting 10 µL of MPXV into the mouse cheek pocket at a concentration of 10^5^ PFU per animal. Intra-anal inoculation was performed by pipetting 10 µL of MPXV into the anal cavity of the mouse at a concentration of 10^5^ PFU per animal.

Following inoculation, mice were weighed every day or every other day and observed for the development of clinical signs of disease and scored against a Human Intervention Point (HIP) matrix. Animals were observed over a time course of 12 or 21 days depending on the study. Tissues were collected at necropsy including nasal turbinate, lung, mediastinal lymph node, heart, kidney, spleen, liver, small intestine, large intestine, brain, bladder, penis and testes which were subsequently analyzed using virological and histopathological methods. Weight was calculated as a percentage of original body weight measured at day 0.

### DNA quantitation from tissues and stool

Viral DNA was extracted from tissues using the Qiagen QIAmp® DNA mini kit (Qiagen, Toronto, Canada Cat# 51306) and from mouse stool using the Qiagen QIAmp® Fast DNA Stool mini kit (Qiagen, Toronto, Canada Cat# 51604) according to the manufacturer’s instructions. For DNA extraction from tissues, samples were incubated at 56°C for 1 hour to completely lyse the tissue and to inactivate MPXV. Stool samples were disrupted in Qiagen InhibitEX® buffer and incubated at 70°C for 5 minutes as per manufacturer’s instructions followed by an additional 1 hour at 56°C to inactivate MPXV. MPXV viral DNA was detected by quantitative PCR using probe and primer sequences and PCR reaction conditions optimized to detect DNA from MPXV Clade II virus as described by Li et al. 2010 ^72^. Specific sequences used were forward primer (5’-CACACCGTCTCTTCCACAGA-3’), reverse primer (5’-GATACAGGTTAATTTCCACATCG-3’) and probe 5’<u>FAM</u>-AACCCGTCGTAACCAGCAATACATTT-3’<u>BHQ1</u> (IDT, Coralville USA). Quantitative probe PCR was performed using the LUNA® Universal Probe qPCR Master Kit (New England Biolabs® Inc., Whitby, Canada Cat# M3004L) Reactions were performed using 4 µL of undiluted DNA, and primers and probes at a final concentration of 0.4 µM each. Reaction annealing temperature was set to 62°C and a total of 45 reaction cycle were used. Remaining reaction parameters followed manufacturer’s instructions. The reactions were performed on a StepOnePlus^TM^ Real-Time PCR System in a 96-well plate (Applied Biosystems, Cat# 4346907) as previously described ^72^.

### Infectious MPXV determination by plaque assay

Tissues were homogenized in serum free MEM using a Qiagen Tissue Lyser with the supernatant of the resulting homogenate used to determine live virus. Plaque assays were performed in 12 well tissue culture plates using BSC-40 cells maintained in culture conditions described above. BSC-40 cells were seeded 24 hours prior to assay at a rate of 2.5×10^5^ cells/well. For each tissue, four supernatant dilutions (5-fold initial dilution and up to 10,000-fold depending on tissue) were assayed in triplicate. A volume of 0.3 mL of supernatant dilutions was added as inoculum to each well and incubated at 37°C in a CO_2_ incubator for one hour. One milliliter of 1% CMC (carboxymethylcellulose supplemented with 2% FBS, 100 U/mL penicillin and 100 ug/mL streptomycin) was added to each well and cells were incubated for 72 hours. CMC was removed, cells were fixed with 10% neutral buffered formalin for 30 minutes and finally stained with 0.4% crystal violet for 30 minutes. Plaque numbers were counted in wells that had 30 or fewer plaques. PFU (plaque forming units) per mg was calculated as total PFU in the volume of homogenate divided by the mass of the tissue homogenized.

MPXV Clade IIb virus stocks were also determined by plaque assays as described above using 0.3 mL of 10-fold dilutions as inoculum.

### Histopathology and Immunohistochemistry (IHC)

Tissues collected for histopathology were placed submersed in 10% buffered formalin (Sigma) for at least 7 days within the CL3 lab prior to being transferred to CL2 facilities for processing. Formalin-fixed tissues were paraffin-embedded, sectioned, slide-mounted, and stained at Prairie Diagnostic Services Inc. (Saskatoon, Saskatchewan). Tissues were stained with Hematoxylin and eosin (H&E).

Immunohistochemical staining was conducted at Prairie Diagnostic Services, Saskatoon, SK using an automated slide stainer (Autostainer Plus, Agilent Technologies Canada Inc., Mississauga, ON). For Vaccinia virus IHC, antigen retrieval consisted of applying Proteinase K (Agilent Technologies Canada Inc., Mississauga, ON) for 10 minutes, followed by a 30-minute incubation of the primary antibody (Rabbit anti-Vaccinia virus, Abcam (catalogue number ab35219) at a dilution of 1:500. For IBA1 antibody (ThermoFisher PA5-27436), antigen retrieval was performed in a Tris/EDTA pH 9 buffer at 97°C for 20 minutes. The primary antibody was used at a 1:4000 dilution and applied for 30 minutes at room temperature. For both, binding of the primary antibody was detected using an HRP-labelled polymer detection reagent (EnVision+ System - HRP Labelled Polymer, Agilent Technologies Canada Inc., Mississauga, ON). The staining was visualized using 3,3’-diaminobenzidine tetrahydrochloride (DAB) (Dako Liquid DAB+ Substrate Chromogen System, Agilent Technologies Canada Inc.) as the chromogen, and counterstained with Mayer’s Hematoxylin.

The stained slides were imaged using a Aperio ScanScope XT (Leica Biosystems, Nußloch, Germany). Visualization and analysis were performed by a board-certified pathologist and given a score ranging from 0 to 4 based on the presence of inflammatory cell types and lesions where a score of 0=absent; 1= slight or questionable; 2= clearly present, but not conspicuously so; 3= marked presence 4= severe.

### Statistical analysis

To assess the effects of time and group on animal weight while accounting for repeated measurements within individuals, a Linear Mixed Model (LMM) was applied in Python (version 3.12) using the statsmodels library (version 0.14.4). The model included time, group, and their interaction as fixed effects. A hierarchical approach was adopted to guide post-hoc testing using Tukey pairwise comparisons. The interaction term (time x group) was examined first; when significant, pairwise comparisons were performed to identify specific differences between combinations of time and group levels. If the interaction was not significant, the main effects of time and group were evaluated independently. Post-hoc pairwise comparisons were then conducted using estimated marginal means with p-values adjusted for multiple comparisons using the Bonferroni correction. Graphs were constructed using GraphPad Prism version 10.0.0 for Windows, GraphPad Software, Boston, Massachusetts USA, www.graphpad.com. A *p* value of ≤ 0.05 was considered statistically significant.

